# Single cone photoreceptors experience global image statistics through gaze shifts

**DOI:** 10.1101/2024.09.07.611793

**Authors:** Takuma Morimoto, Luna Wang, Kinjiro Amano, David H. Foster, Sérgio M. C. Nascimento

## Abstract

Our visual experience does not merely reflect a static view of the world but is a dynamic consequence of our actions, most notably our continuously shifting gaze. These shifts determine the spectral diet of any individual cone photoreceptor. The aim of this study was to characterize that diet and its relationship to scene adaptation. Gaze shifts were recorded from observers freely viewing scenes outdoors for five minutes. Hyperspectral images of the scenes were also recorded from the observer’s eye position. As a control, gaze shifts were also recorded from observers viewing the images on a computer-controlled display in the laboratory. From the hyperspectral data, spatially local histograms of estimated excitations in long-, medium-, and short-wavelength-sensitive cones were accumulated over time at different retinal locations. A global illuminant change was then introduced to test how well local retinal adaptation discounted its effects. The results suggest that over short periods individual cones tend to experience the statistics of full scenes, with local adaptation compensating for illumination changes almost as well as global adaptation. This compensation may help to maintain our stable local perception of scene colour despite changes in scene illumination.

## 1. Introduction

Characterizing our everyday exposure to natural optical environments may offer insights into the role of individual visual mechanisms. These mechanisms need to deal with the complex varying patterns of spectral radiation incident at the eye [1-3], sometimes referred to as our ‘spectral diet’ [4]. Yet quantifying the exact signals received by individual photoreceptors presents a challenge: our spectral diet is not a passive reflection of a static external world, but is dynamically shaped by our active behaviours, most importantly our shifting gaze. There have been efforts to characterize physical regularities in natural scenes and their connection to a range of visual functions (e.g. [5-13]), and some studies have considered how our sampling behaviour interacts with experienced image statistics [14-20].

The problem of concern here is this. At any given moment, alongside the dynamics of our behaviour, the complex spatial structures of natural environments [21,22] provide different physical inputs to cones at different retinal positions. This suggests that our cones, distributed over the retina, receive markedly different physical inputs from moment to moment, potentially affecting visual function across the visual field. In some species, there are structural consequences. An extreme example is the visual field of mice, where only the upper visual field appears to support colour discrimination [23], presumably in response to the asymmetry between mouse views of the ground and the sky [24]. With our less stereotyped behaviour, we can counter the spatial inconsistency of photoreceptor experience by shifts in gaze, which ensure each cone experiences a variety of spectra over time, cumulatively more representative of the entire scene. The first goal of this study was to formally test the extent to which our natural sampling behaviours enable individual cones to access global scene statistics.

Characterizing local spectral diets is particularly relevant in colour vision research because retinal adaptation is thought to be one of the main mechanisms for colour constancy [25-28], a visual ability to perceive object or surface colour consistently despite changes in the spectrum of the illumination. The required adaptation is routinely achieved by scaling each class of cone signals across the whole retina by a single multiplicative factor [29]. This operation can be represented mathematically by a diagonal matrix transformation [29], which has been the basis of many computational colour constancy algorithms [30] and colour appearance models, standardized by the Commission Internationale de l’Eclairage (CIE), such as CIECAM02 and CIECAM16 [31, 32]. Yet von Kries originally proposed this scaling as a local, not global, mechanism [27, 33]. Local scaling may be more plausible at the photoreceptor level but its efficacy seems not to have been empirically tested. The second goal of the present study was to assess how well local retinal adaptation approximates the effects of global retinal adaptation.

To achieve these goals, both observer gaze data and spectral radiance data are needed at each point in a scene to estimate the spatial pattern of cone excitations across a simulated retina. Although gaze data with RGB images are widely available [34-37], gaze data with hyperspectral images for radiance calculations are less accessible. Accordingly, gaze patterns were recorded in seven outdoor environments with observers freely viewing each scene for five minutes. Hyperspectral radiance images of each scene were also acquired from the observer’s eye position. In computational simulations with these data, the history of individual cone excitations was then estimated at different retinal locations. It was found that even over short periods, individual cones tend to experience the full scene statistics and that the resulting local adaptation can compensate for scene illumination changes almost as effectively as global adaptation.

## 2. Methods

Gaze recordings and hyperspectral images were obtained from seven different outdoor scenes. Two experienced observers viewed the scenes outdoors and, as a control, the same observers and three other naïve observers viewed the scenes on a computer-controlled display in the laboratory. In both conditions, the emphasis was on capturing natural behaviours in free viewing, without the influence of task demands such as searching for or detecting a specific target.

### (a) Gaze tracking

Gaze patterns were recorded from the observer’s right eye with an infrared eye tracker system (iView, version 3.01, SensoMotoric Instruments, Germany), attached to a chin and forehead rest anchored to a table, as illustrated in Figure 1a. Eye position was estimated from the corneal reflection. The system had a sampling rate of 50 Hz (i.e. data interval 20 ms) and the tracking area in the frontoparallel plane subtended a visual angle 30° horizontally and 25° vertically. Measurements outdoors were made under a parasol to prevent direct light from reaching the eye tracker (Figure 1b). For the laboratory measurement (Figure 1c), observers were positioned in front of a CRT monitor (GDM-F400T9, 19-inches, 1600×1200 pixels, Sony, Tokyo, Japan) with the eye tracker system. Scene images were displayed on the monitor, with the same visual angle as the outdoor measurement, through a graphics card (VSG 5, Cambridge Research Systems, Rochester, UK) that allowed 8-bit depth per each RGB phosphor. The range of pixels outside of the monitor’s chromaticity gamut varied from 0.0% to 6.9% across 7 scenes. Overall luminance levels were reduced to fit the luminance range allowed by the monitor, and the mean luminance level ranged between 8 and 15 cd/m^2^.

**Figure 1.**
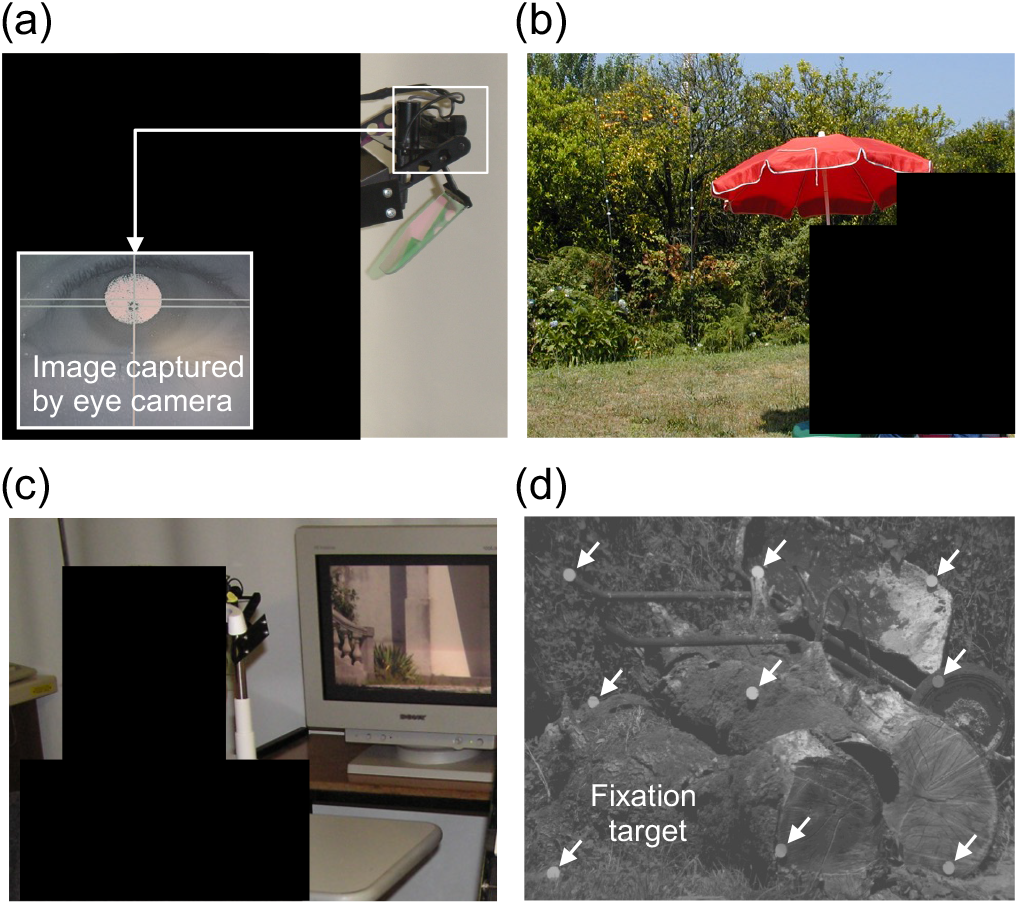
(a) The system used for the measurement of gaze position. (b) Gaze measurements made outdoors. (c) Gaze measurement in the indoor laboratory environment. (d) Nine fixation targets placed in a scene to calibrate the eye tracker. Black squares have been added for privacy reasons.

For calibration, nine grey fixation targets were inserted into the scene to form an approximately regular array from the observer’s viewpoint. A grayscale image with the fixation marks in place (Figure 1d) was loaded into the eye-movement software. The observer looked at each mark in turn during the calibration procedure.

The raw gaze data contained both fixations and saccades, but all were included in the analysis, which was concerned solely with cone spectral diet over time.

### (b) Observers

For outdoor measurements, the observers were authors KA and DF, both male, aged 33 and 59 years, respectively. For indoor measurements, these observers were joined by three other observers (EO, MA and NM), also male, aged 24–34 years. All observers had normal or corrected-to-normal visual acuity and normal colour vision as assessed with an anomaloscope (Oculus HMC Anomaloscope, Oculus, Wetzlar, Germany).

### (c) Hyperspectral imaging

The seven hyperspectral radiance images were acquired as described previously [38, 39], with the hyperspectral camera placed approximately at the observer’s viewpoint. Colour images are shown later. Where necessary for computational purposes, spectral radiance images were converted to effective spectral reflectance images by dividing pointwise by the estimated spectrum of the direct illumination recorded from a reference surface in the scene [40]. To supplement the main analysis, an additional 45 hyperspectral reflectance images were also used [41]. Each hyperspectral image had dimensions approximately 1344 × 1024 pixels, corresponding to a camera angle of approximately 6.9° × 5.3° and spectral range 400–720 nm sampled at 10 nm intervals. To reduce non-imaging noise in the unaveraged source data and to shorten computational time, the spatial resolution of all images was adjusted to a size of 336 × 256 pixels by spatial averaging [13].

### (d) Viewing procedure

Each observer, after completing the gaze calibration, freely viewed the scene for 5 minutes, in the knowledge that they would be asked a randomly chosen question about the contents of the scene (e.g. how many tree trunks were present). A fixation recalibration (recentering) was carried out every minute during the viewing period. At the end of the period, the gaze calibration was checked, and, if valid, the gaze data were saved. For one observer, there was a small gaze misalignment with two of the seven scenes, which was not improved after repeating the observation.

Experimental procedures complied with current guidelines of the Research Ethics Committee of the University of Minho to the Colour Science Laboratory (CEICVS 052/2021).

## 3. Analysis of gaze patterns

Figure 2 shows a schematic of the analysis. On the left, the “Observation of a scene” image represents an observer shifting their gaze at hypothetical times *t*_1_ to *t*_4_. On the right, the “Spatial filtering” image depicts the inverted retinal image within a 2.5°-radius region at time *t*_1_. This angular range was chosen to fit within the acceptance angle of the hyperspectral imaging system and corresponds approximately to the size of the fovea [42-43]. To accommodate varying cone density with eccentricity, the projected image was convolved with a two-dimensional Gaussian filter with standard deviation 1 pixel at the central fovea, corresponding approximately to 1 arcmin, the minimum angle of resolution. As eccentricity increased away from the central fovea, the standard deviation was increased by the reciprocal of the cone density [44], depicted below the circular image. The result is shown in the “Retinal Images” *f*_1_, *f*_2_, *f*_3_, *f*_4_ at times *t*_1_, *t*_2_, *t*_3_, *t*_4_, respectively.

**Figure 2.**
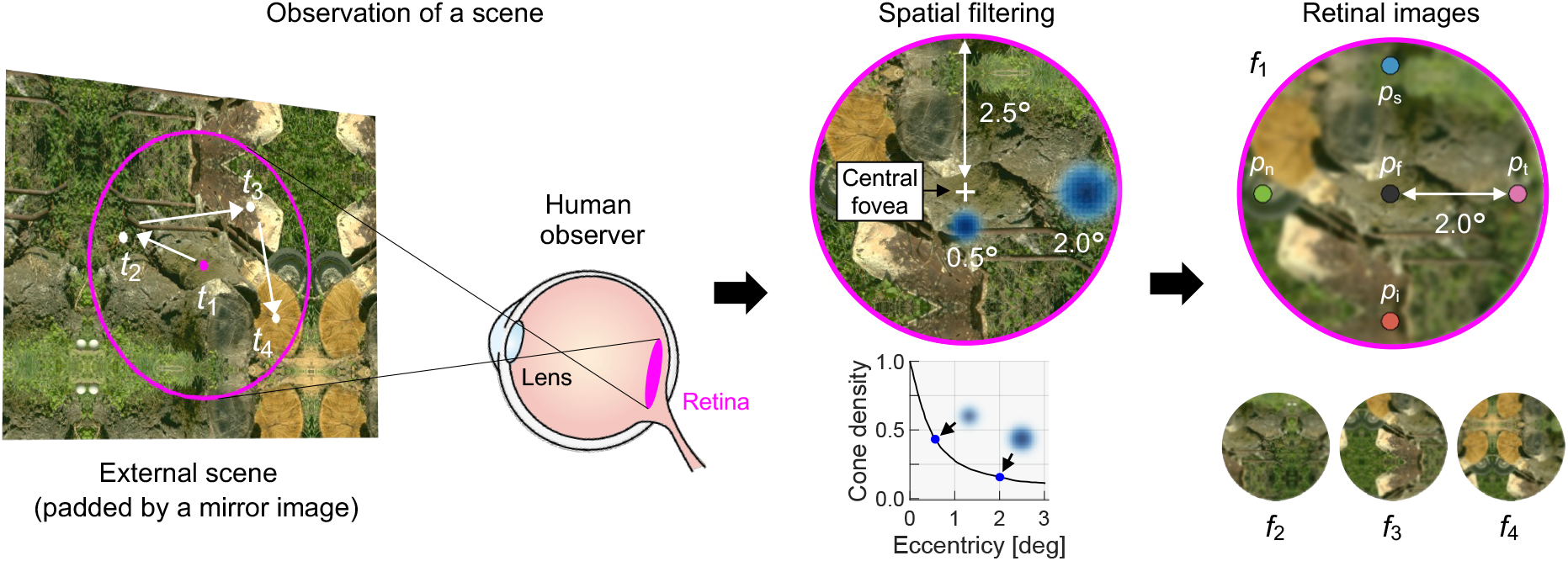
Schematics of the simulation and analysis using recorded gaze data. At a given time point *t*_1_, a circular portion of the scene is projected onto the observer’s retina. Spatial filtering is applied using a 2D Gaussian, with the standard deviation determined by cone density at the corresponding eccentricity. This produces retinal frames *f*_1_, *f*_2_, *f*_3_ and *f*_4_ sampled at time *t*_1_, *t*_2_, *t*_3_, and *t*_4_, respectively. In subsequent analyses, photoreceptors at five locations are considered: the fovea *p*_f_, inferior retina *p*_i_, superior retina *p*_s_, and nasal retina *p*_n_ and temporal retina *p*_t_.

Possible systematic differences in spectral diet were tested at five different retinal locations: the centre of the fovea *p*_f_ (black filled circle), inferior retina *p*_i_ (red filled circle), superior retina *p*_s_ (blue filled circle), nasal retina *p*_n_ (green filled circle), and temporal retina *p*_t_ (pink filled circle). Each of these five test locations corresponds to a single pixel in the filtered retinal image. The hyperspectral image was padded with its own mirror image to prevent edge discontinuities and to avoid the central test location sampling more points than the other locations. The imaged area of the original scene was thus assumed to be characteristic of the unimaged areas.

In the following, the spatial scale of each of the five test locations is referred to as local and of the whole image as global.

### (a) History of cone excitations at different retinal locations

The relationship between local and global photoreceptor sampling was quantified by the correlation between local and global histograms of cone excitations. Figure 3 illustrates at each retinal location the time-course of cone excitations, based on the 2-degree Stockman and Sharpe cone fundamentals [45, 46]. The square coloured patches show RGB renderings of spectra sampled at two retinal locations *p*_s_ and *p*_t_ for the four retinal images or frames *f*_1_, *f*_2_, *f*_3_, *f*_4_ sampled at times *t*_1_, *t*_2_, *t*_3_, *t*_4_, respectively. Below are shown corresponding histograms of the logarithm to the base 10 of L, M, S cone excitations accumulated over time from the start of the measurement with the first image (first frame) to a specific end frame *f*_*n*_ (*f*_4_ in this example), where *n* increased from 50 to 15,000 in increments of 1. The top two rows of histograms are for local excitations and the bottom row for global excitations.

**Figure 3.**
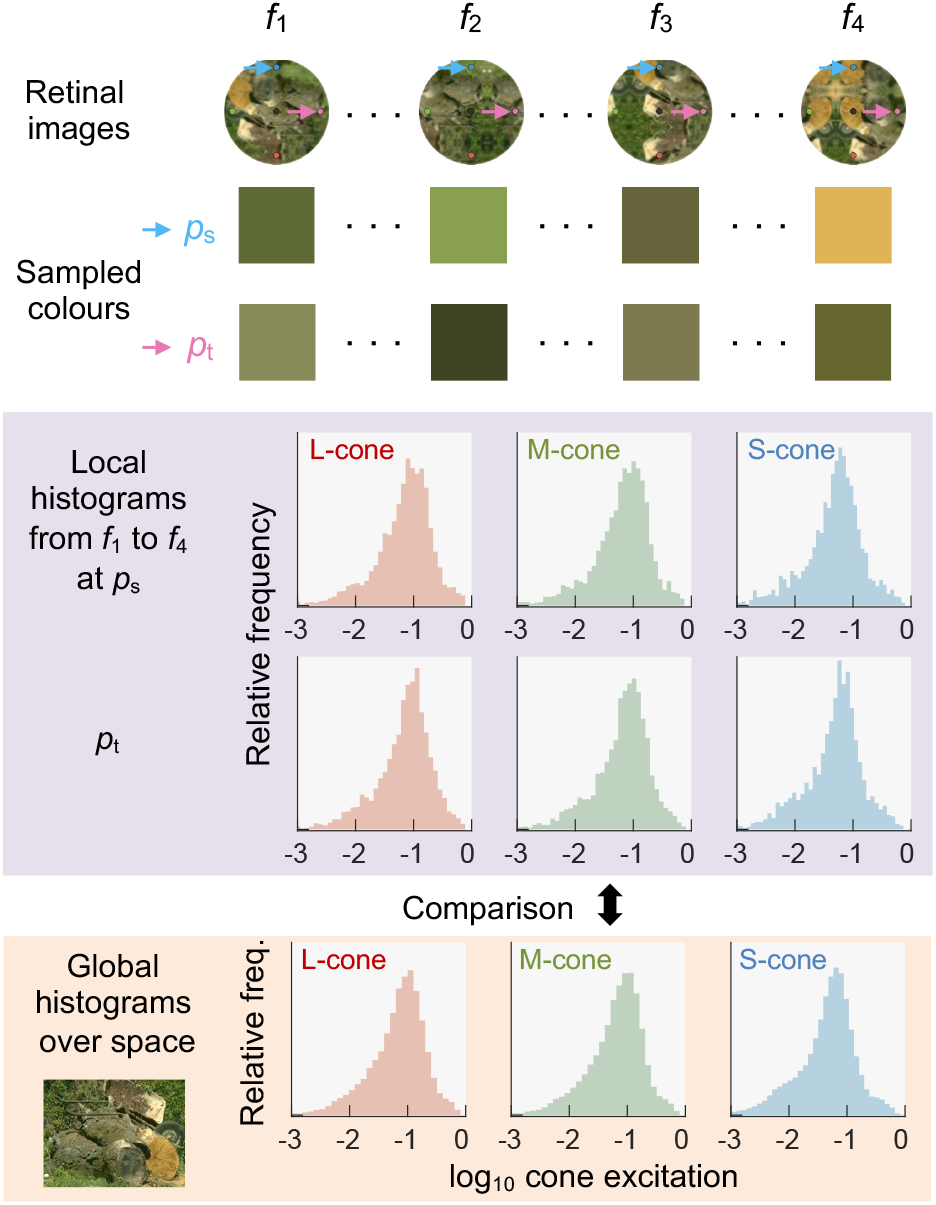
History of excitations of individual cones at different retinal locations. The top four circular images display retinal images *f*_1_, *f*_2_, *f*_3_ and *f*_4_, with RGB renderings of spectra sampled at *p*_s_ and *p*_t_ shown below. The top two rows of histograms represent local excitations accumulated from *f*_1_ to *f*_4_ at *p*_s_ and *p*_t_, respectively. The bottom row shows the global histogram drawn from the entire image.

The number of data points in the local histograms depended on the end frame. For instance, the first histogram used 50 data points and the last histogram used 15,000 data points. The bin number was systematically determined by the Freedman-Diaconis rule [47], which is robust to outliers. The same bin number was used for global histograms from the whole image. The correlation between the relative frequencies of the global and local histograms was obtained separately for each retinal location, and repeated for all end frames, resulting in 14,951 product-moment correlation coefficients. At the centre of the fovea, S-cone excitations were omitted owing to the sparsity of S cones there [48].

The extent to which samples from individual cones capture global scene statistics should be evident in an increase in correlation coefficient between local and global histograms with time from gaze onset. For indoor gaze measurement, this analysis was conducted only for the centre of the fovea (*p*_f_) because padding could not be used due to the absence of visual stimuli beyond the experimental monitor.

### (b) Local retinal adaptation and discounting the influence of illumination

The extent to which local retinal adaptation discounts the effects of a global illumination change was quantified colorimetrically. Recall that in normal viewing conditions, retinal adaptation to a change in illumination is rarely complete [49, 50], and colorimetric models of chromatic adaptation usually incorporate a weighting coefficient, the degree of adaptation *D*, which ranges from 0 to 1 [51]. For simplicity, this factor *D* is omitted from the present calculations.

The local adaptation was estimated at time *t*_*n*_ from a weighted sum of the history of retinal images from the first frame *f*_1_ until *f*_*n*_. As illustrated by the plot of weight against time in Figure 4, weights decreased exponentially from the largest associated with the most recent image. The estimated adaptation was applied pixel-wise in a diagonal matrix transformation to the retinal image sampled at the subsequent time *t*_n+1_. Because of computational demands, just six discrete times were used with roughly equal logarithmic steps from *t*_3_ to *t*_14,999_, where *t*_3_ was chosen as the start since there is evidence that an exposure of 60 ms can influence subsequently perceived colour [52]. If observers uniformly sampled an image, the expectation would be that all cones would adapt to the scene’s mean colour and local adaptation would coincide with global adaptation.

**Figure 4.**
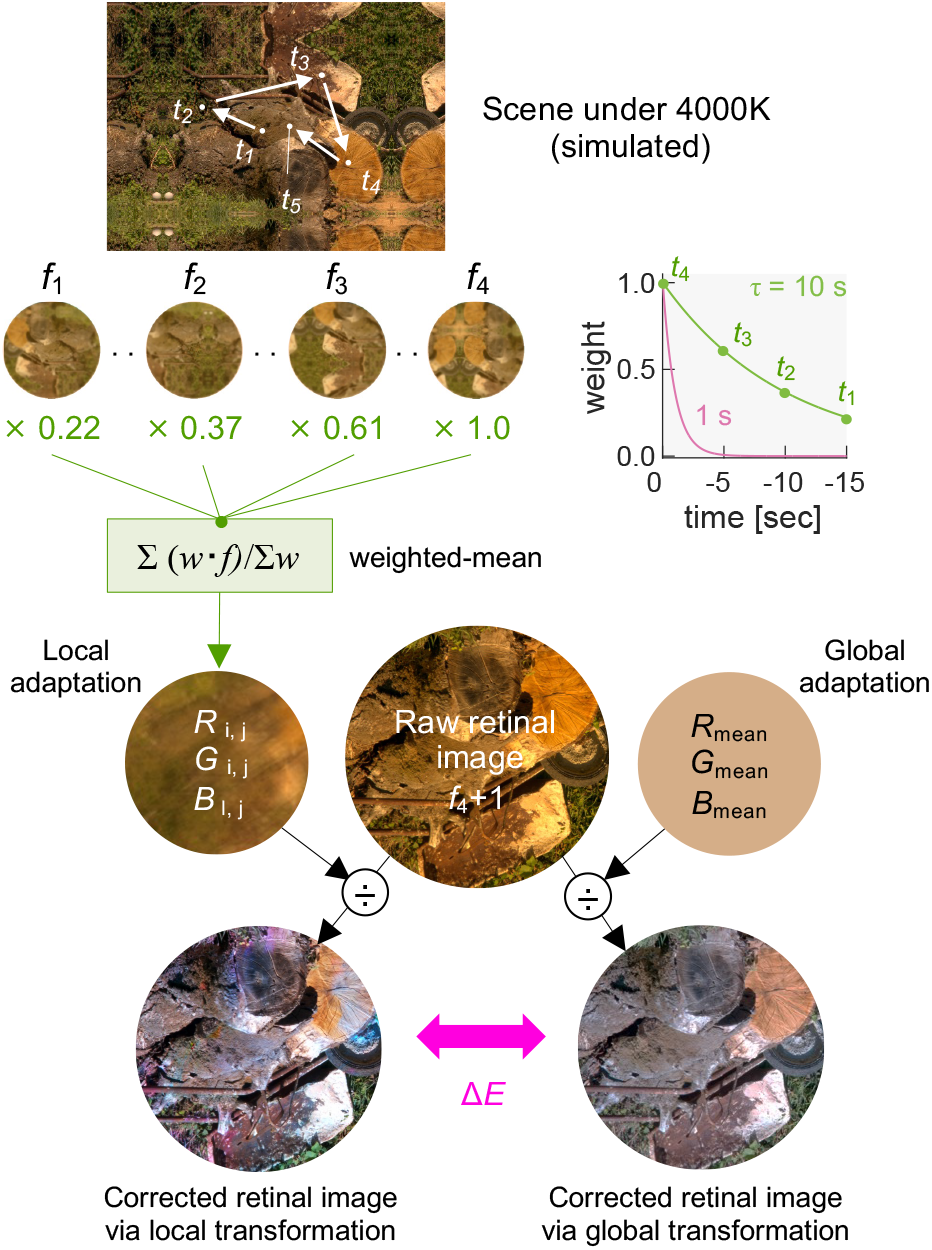
Effect of local adaptation on discounting a global illuminant. The local adaptation state was estimated from a weighted sum of past retinal images, while global adaptation relied on the mean color of the entire scene. After illuminant correction, color differences Δ*E* between images corrected for local and global adaptation were evaluated.

The mean correlated colour of the direct illumination recorded from the reference surface in each scene was 5,571 K with range 4,421 K to 7,643 K. To provide a controlled change in illuminant, the effective spectral reflectance image of the scene was multiplied pointwise by a daylight illuminant with CCT 4,000 K [53], which approaches the lower limit of outdoor illuminant changes [54] and sufficient here to test chromatic adaptation.

Colour differences Δ*E* between images corrected for local and global adaptation were evaluated in an approximately uniform colour appearance space CIECAM16-UCS [55], which uses the CIE 1931 cone fundamentals rather than those of Stockman and Sharpe [45, 46]. This space is more uniform than CIELAB colour space [53], also used in similar analyses. Spectral radiances at each point *i, j* under a 4,000 K global illuminant were converted to tristimulus values *X*_*i,j*_,*Y*_*i,j*_,*Z*_*i,j*_ and then to cone-like responses *R*_*i,j*_,*G*_*i,j*_,*B*_*i,j*_ defined in CIECAM16-UCS thus:

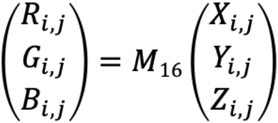

where

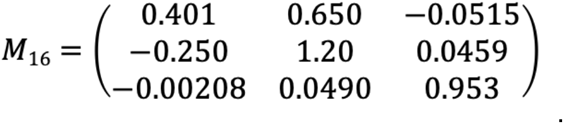

After local diagonal matrix transformation, the corrected cone excitations are given by

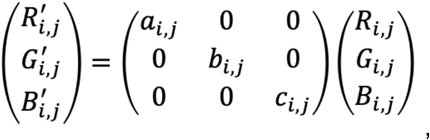

where the scalars *a*_*i,j*_, *b*_*i,j*_, *c*_*i,j*_ depend on spatial position *i, j* and are set by the reciprocal values of *R*_*i,j*_, *G*_*i,j*_, *B*_*i,j*_ in the locally adapted retinal image. Analogously, after the global diagonal matrix transformation, the corrected cone excitations are given by

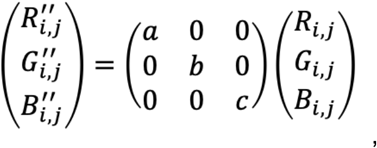

where the scalars *a, b, c* are constants that depend only on the global spatial means. Note that the automatic chromatic adaptation stage in CIECAM16 was omitted to avoid double adaptation.

Figure 4 illustrates a hypothetical sequence where the observer shifts gaze at time *t*_1_ to time *t*_5_ (top image), where *t*_4_ is the current time and *t*_5_ is the time of the next frame, 20 ms after *t*_4_. Estimates for two time constants, 1 s and 10 s, are shown, which fall within the range reported previously [56-60]. Local and global diagonal matrix transformations were applied to discount the illuminant influence (bottom images).

Colour differences Δ*E*_*i,j*_ between the local and global images were calculated at each point *i, j*, and the median taken to avoid the influence of outliers. A value of zero at every point indicates the same cone adaptation locally and globally and a positive value indicates spatial inhomogeneity. Although gaze patterns were not recorded with images of scenes under a 4,000 K illuminant, it was assumed that gaze patterns were the same as with the original images.

### (c) Simulated observers

To compare with human behaviours, two model observers were simulated: one used random gaze, where gaze position was chosen randomly in each frame; the other used random walk, where gaze position was the result of a random shift in direction with the fixed step size [61, 62]. For both models, gaze positions were constrained to fall within the image region. Step size for the random walk model were based on recorded angular velocities, with values equal to the mode (2.3 deg s^-1^) and one-third and three times the mode (0.77 deg s^-1^ and 6.9 deg s^-1^). Gaze positions of these simulated observers were calculated for 5 min at the same temporal frequency as with the eye tracker, resulting in 15,000 frames. The gaze position with the first frame was set to the same random value for both models and their variations. Data were analyzed in the same way as with human gaze.

## 3. Results and comment

### (a) Gaze shifts in each scene

Figure 5 displays images of the scenes, with the corresponding gaze patterns measured outdoors for two of the five observers, DF and KA, shown on the right of each image.

**Figure 5.**
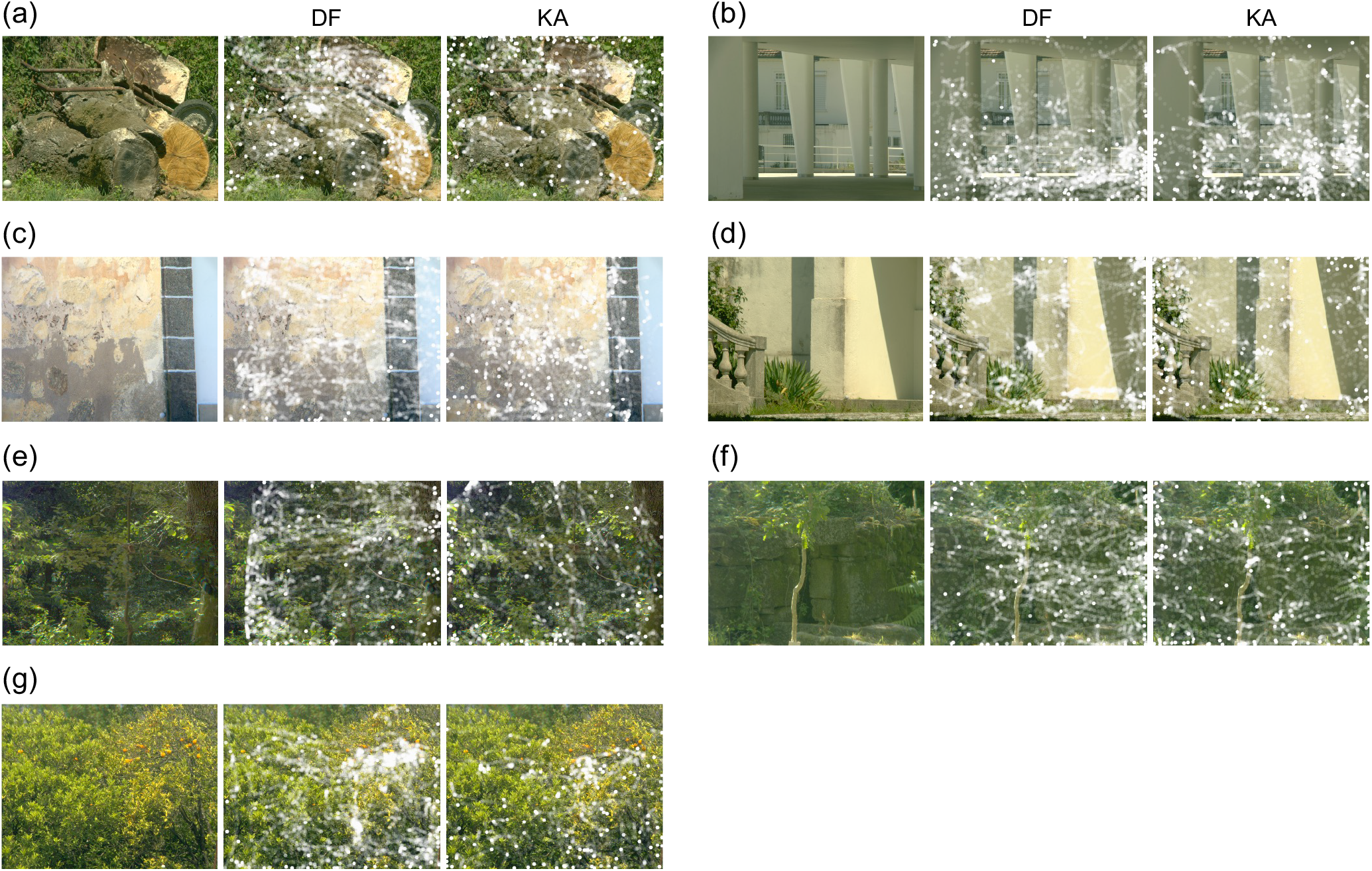
Images of seven natural scenes and to the right of each the gaze patterns measured outdoor for two observers.

### (b) Correlation between sampling by individual cones and global scene statistics

Figure 6 summarizes the time course of the correlations in the outdoor gaze measurements. The first three columns show trajectories of the product moment correlation coefficient between local and global histograms for L, M and S cone classes and observers DF and KA. Retinal locations and their color codes are included in the upper plot of panel (a). The correlation increased over time in a similar way across cone classes. For observer DF, it increased rapidly during the first 10 s and gradually reached a maximum value between 0.75 and 0.89. For observer KA, it increased more gradually, but eventually a maximum between 0.66 and 0.78.

**Figure 6.**
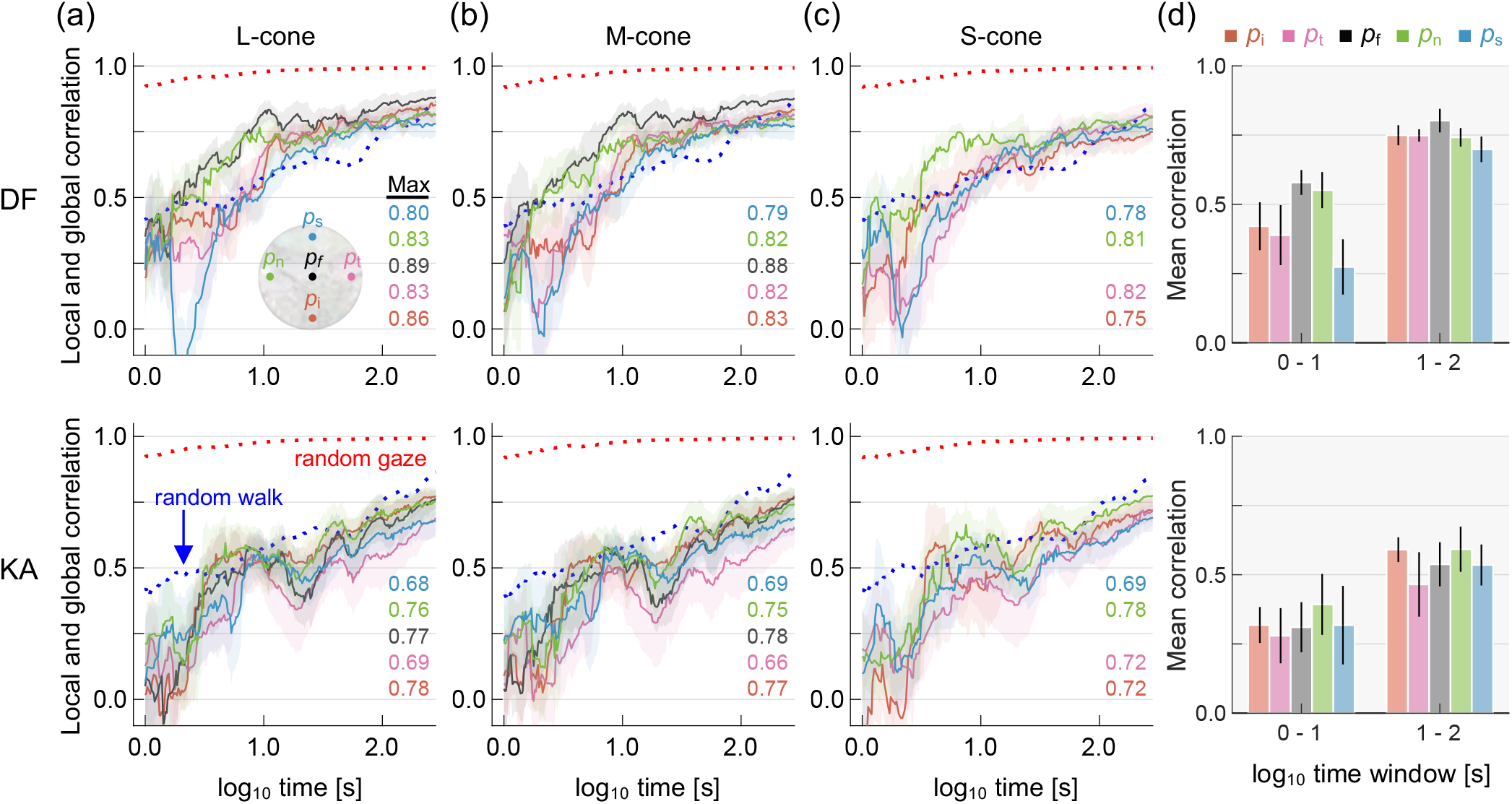
Correlation between local and global statistics over time for outdoor gaze measurements. The first three columns show the product moment correlation coefficient between local and global histograms for each cone class averaged over seven scenes plotted against the base 10 logarithm of the time from gaze onset. The two rows show data for observers DF and KA. The colours of the curves indicate retinal location and the shaded area represents ±1 standard error (SE) across scenes. The dotted red and blue curves represent predictions of random gaze and random walk models, respectively. The last column shows the mean product moment correlation coefficient over two time windows 1-10 s and 10-100 s and the colours of the bars indicate retinal location (upper panel in panel a). Error bars represents ±1 SE across scenes.

Predictions from the random gaze model and random walk model with mode angular velocities are shown by red and blue dotted lines, respectively, averaged across 5 iterations. The random gaze model provides a much poorer fit than the random walk model, presumably because with random gaze, locations are unrelated and may be far from each other, whereas with random walk, locations change progressively, approximating the behaviour of observers exploring a scene. The panel (d) in Figure 6 shows the correlation coefficient averaged over L and M cones, seven scenes, and two different time windows (1–10 s and 10–100 s). Data for S cones were omitted at *p*_f_.

From the data shown in panel (d), a two-way analysis of variance (ANOVA) for retinal location (5 levels) and time windows (1–10 s and 10–100 s) for each observer revealed a significant main effect of time window (*F*(1,6) = 83.8, *p* < 0.001 for DF; *F*(1,6) = 10.9, *p* < 0.02 for KA), but no significant main effect was found for retinal locations (*F*(4,24) = 2.2, *p* = 0.09 for DF; *F*(4,24) = 0.52, *p* = 0.7 for KA). Also, the interaction between the two factors was not significant (*F*(4,24) = 1.4, *p* = 0.26 for DF; *F*(4,24) = 0.15, *p* = 0.96 for KA).s

As shown in Figure 7, the corresponding correlations were similar in the indoor gaze measurements with five observers. The S-cone data are omitted as the analysis concerns only the centre of the fovea *p*_f_. For all observers, the correlation coefficient rapidly increased during the first 10 s and gradually increased thereafter. The maximum correlation ranges from 0.87 to 0.94, generally showing higher values than the outdoor measurement. The trend is similar for L and M cones.

**Figure 7.**
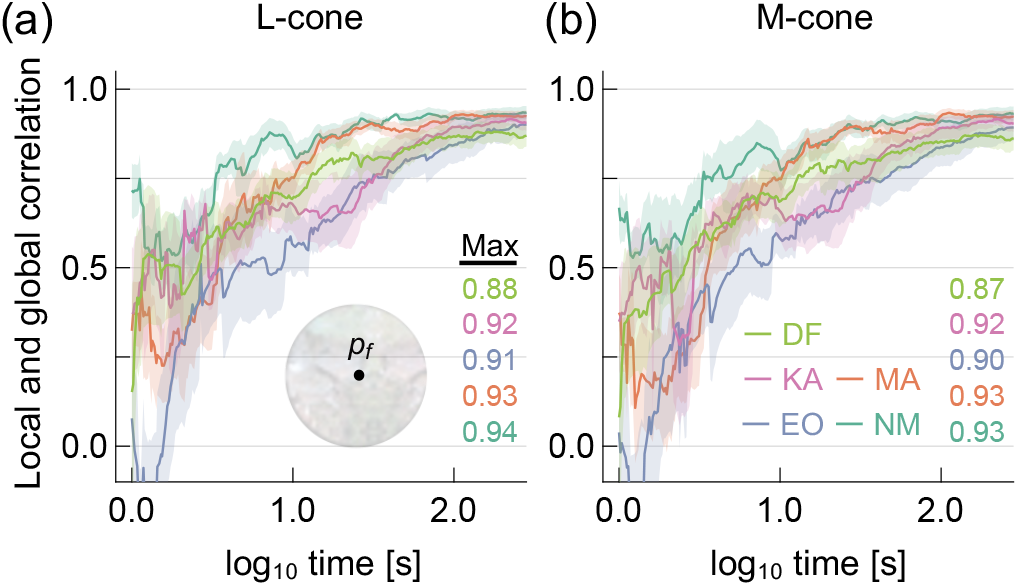
Transition of correlation coefficient between local and global statistics over time for indoor measurements for (a) L cones and (b) M cones. Data for S cones were omitted as the analysis here is for the centre of the fovea *p*_f_. Observers are indicated by different colours. DF and KA also participated in the outdoor measurements. The shaded area represents ±1 standard error (SE) across scenes.

Taken together, these findings suggest that over time excitations at individual cones approximate global scene statistics, irrespective of photoreceptor locations.

### (c) Local retinal adaptation and von Kries’ scaling

Figure 8 shows the time course of the median CIECAM16-UCS colour difference Δ*E* between images corrected by local and global diagonal matrix transformations. The two columns refer to two different adaptation time constants.

**Figure 8.**
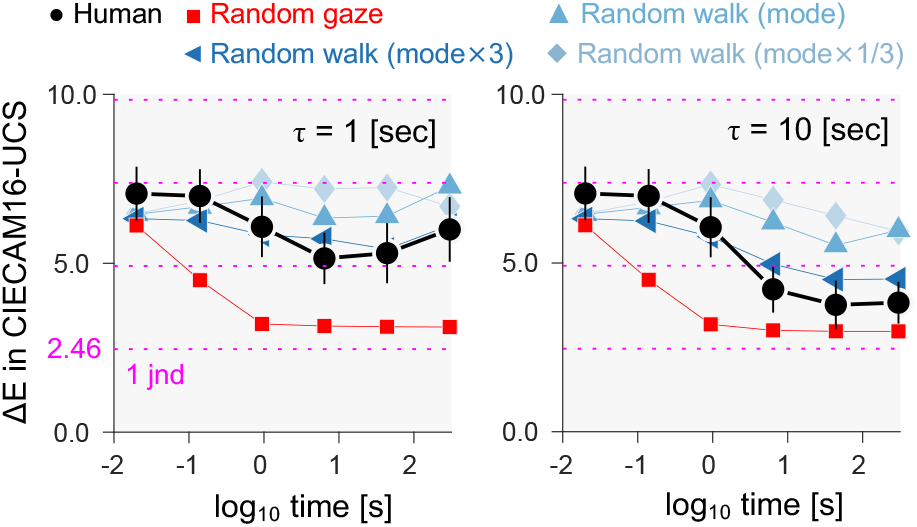
Discounting illumination colour by local retinal adaptation over time. Median CIECAM16-UCS colour differences Δ*E* between images corrected by local and global diagonal matrix transformations are plotted against the logarithm of the time from gaze onset. Data are averaged across two observers and seven scenes, with simulations based on random gaze and random walk. The horizontal dotted line shows increments in multiples of 1 jnd.

For detecting colour differences Δ*E* in images of natural scenes, a threshold value or just noticeable difference (jnd) in CIELAB space has been estimated as about 2.2 for images of the kind used here [63]. Although there is no unique conversion between CIELAB thresholds and CIECAM16-UCS thresholds, separate simulations with the 52 images in this study suggest that a CIELAB colour difference of 2.2 corresponds approximately to a CIECAM16-UCS colour difference of 2.5, which was accordingly used as the jnd. This approach is similar to that used previously in transforming jnd values between colour spaces [41].

Overall, color differences were slightly smaller for a time constant of 10 s than for 1 s, as expected. To confirm this statistically, minimum Δ*E* values were calculated for each of the 7 scenes at both time constants (1 s and 10 s). A one-way repeated measures ANOVA revealed a significant effect of time constant (*F*(1,6) = 46, *p* < 0.001) on the minimum Δ*E* averaged across scenes. The influence of the 4,000 K illuminant was significant, evidenced by the fact that without an illuminant correction the colour difference Δ*E* was 29 averaged across scenes. Against this value, the local diagonal matrix transformation based on the first three image frames brought marked benefit, with Δ*E* values decreasing progressively to a minimum around the two end time points (44 s and 300 s). Note that even with a shorter time constant of 1 s, the lowest Δ*E* was 4.8, about 2 jnd value*s*.

For scenes where pixel chromaticities varied little over space and the mean chromaticity was close to grey (e.g. scenes in panels (b) and (c) in Figure 5), Δ*E* values were small at the first frame but then decreased more slowly afterwards, showing that the benefits of gaze shifts depend on chromatic complexity. As before, the random gaze model provided a much poorer fit than the random walk model.

The closeness of the random walk model to observer performance allows a test of the generalizability of the present findings from seven natural scenes to larger samples, namely the additional 45 images shown in Figure 9. As the plots in Figure 9 show, the time courses of the colour differences Δ*E* are similar, though the minima are slightly higher than in Figure 8.

**Figure 9.**
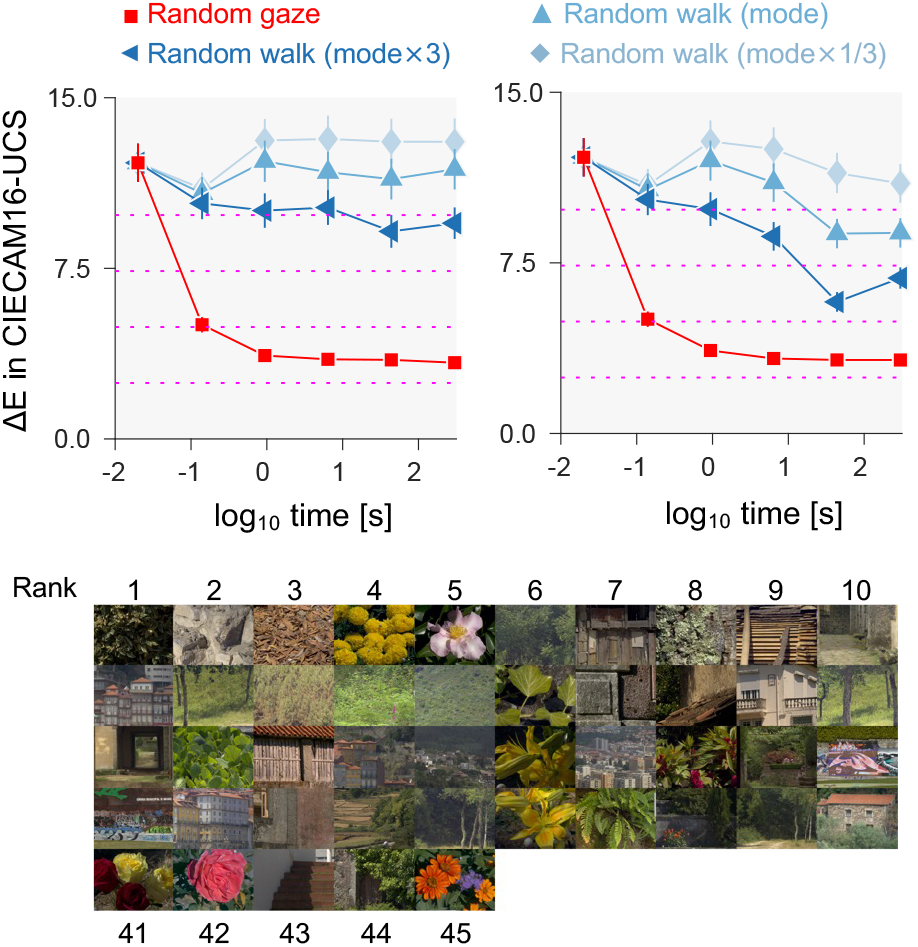
Simulations with random gaze and random walk averaged across the 45 scenes in the montage. Images are shown in ascending order of minimum Δ*E* for random walk model (three-times mode). The horizontal dotted line shows increments in multiples of 1 jnd.

In summary, it seems that gaze shifts can indeed discount the adaptational effects of global illumination changes, especially over longer time periods [64]. These trends hold over large samples of scene images.

## 4. Discussion

We routinely encounter a variety of natural optical environments, often with complex structures. How single cones accommodate this changing spectral diet was addressed here by analysing gaze patterns in relation to the spectral structures of scenes. Two main mechanisms were identified. One was the convergence over time between the physical signals experienced by individual cones and global scene statistics, regardless of retinal locations. This implies that our active sampling behaviour largely standardizes the spectral diet of individual cones, at least within the foveal region. The other mechanism was the convergence over time between the effects of local adaptation in discounting illumination changes and global adaptation based on average scene colour. This points to the importance of local gain control in shaping colour constancy under varying lighting conditions.

### (a) Adaptation to visual environments

As noted earlier, visual adaptation can occur over multiple time scales [59, 65-68], and a variety of studies have demonstrated the normalizing effect of chromatic adaptation. These include eliminating colour differences between fovea and periphery by adapting to macular filtering [69]; changes in colour perception after wearing coloured glasses [70]; violet-blue colour shifts diminishing after cataract surgery [71]; and seasonal biases in wavelength settings of unique yellow [72]. The present analysis suggests that adaptation by gaze shifts can be added to this resource of normalizing mechanisms.

### (b) Implications for colour constancy

Although our spectral diet is known to play an important role in colour appearance and colour constancy [73, 74], the relationship between local and global mechanisms has received less attention [25]. The sampling of the larger environment identified here is crucial. Without the spatial averaging delivered by eye movements, shadowed regions in a scene might look more like directly illuminated regions, an effect analogous, at a different scale, to the fading experienced in Troxler’s effect.

### (c) Limitations

This study had several technical constraints, which should not affect the conclusions. First, the hyperspectral imaging system had a limited acceptance angle of about 6 degrees, which reduced angle-dependent variations in spectral tuning. In the absence of outdoor measurements, this limitation could have been circumvented in the laboratory by reducing the distance between observer and computer-controlled display Second, the fifty-two hyperspectral images used in the study were not fully representative of real-world scenes. Most, for instance, included little or no sky, which by contrast with the ground would have introduced large chromatic and luminance variation across the retina [75]. Third, although a simple decaying exponential model of adaptation sufficed, it could have been extended to include higher-order temporal filtering [76], together with photoreceptor noise [77] and the detailed variation in the retinal cone mosaic [78, 79]. Finally, gaze data were gathered for a free-viewing task, because of the difficulties with implementing a random target search task outdoors. The nature of the task and visual attention are both known to influence gaze behaviour [80-82]. Coincidentally, free viewing may have contributed to the success of the random-walk model in approximating observer responses, given its indifference to scene content.

## 5. Conclusion

This study showed how the spectral diet of cones is mediated by natural shifts of gaze in outdoor environments. In the course of sampling scenes, individual cones may experience close to the full scene statistics, so that the resulting local adaptation compensates for scene illumination changes almost as well as global adaptation. This compensation may help to maintain a stable local perception of scene colours despite natural changes in scene illumination.

## Declaration of interests

The authors declare no competing interests.

## Acknowledgements

TM was supported by a Sir Henry Wellcome Postdoctoral Fellowship from Wellcome Trust (218657/Z/19/Z) and a Junior Research Fellowship from Pembroke College, University of Oxford. This work was supported by the Portuguese Foundation for Science and Technology (FCT) in the framework of the Strategic Funding (UIDB/04650/2020), the Leverhulme Trust (RPG-2022-266), and the EPSRC (EP/W033968/1). For the purpose of open access, the author has applied a CC BY public copyright license to any Author Accepted Manuscript version arising from this submission.

## Notes

### Competing Interest Statement

The authors have declared no competing interest.

